# Opinions of 12 to 13-year-olds in Austria and Australia on the worry, cause and imminence of Climate Change

**DOI:** 10.1101/333237

**Authors:** Inez Harker-Schuch, Frank Mills, Steven Lade, Rebecca Colvin

## Abstract

Although we are in the third decade of climate science communication as a discipline, and there is overwhelming scientific consensus and physical evidence for climate change, the general public continues to wrestle with climate change policy and advocacy. Early adolescence (12 to 13 years old) is a critical but under-researched demographic for the formation of attitudes related to climate change. This paper presents opinions on the worry, cause, and imminence of climate change that were collected from *n*=463 1^st^ year secondary school students (12-13 years old) in public secondary schools in inner-urban centres in Austria and Australia. Overall, 86.83% of eligible respondents agreed that climate change was probably or definitely something we should worry about, 80.33% agreed that climate change was probably or definitely caused by humans, and 83.17% agreed that climate change was probably or definitely something that was happening now. The respondents’ opinions were also compared to their respective adult population, with Australian 12-13 year olds showing strong positive climate-friendly attitudes, both in comparison to their adult population, and to their Austrian peers. In addition, although the opinions of Austrian 12-13 year olds were quite high, they did not reflect the higher climate-friendly opinions of their adult community. Our results suggest that socio-cultural worldview or socio-cultural cognition theory may not have the influence on this age group as it does on the respective adult population – and, if they are affected, there are attitudes or factors in this age group which resist the opinion-influence from their mature community. These findings are significant as early adolescents may be pivotal in the climate science communication arena and investigating their opinions with regard to climate change may offer an unexplored and under-utilised target for future communication efforts and climate literacy programmes.

## Introduction

Despite more than 30 years at the forefront of the political and social agenda, meaningful climate change governance continues to be thwarted by disconnects between scientific knowledge, public knowledge and trust of climate science. A great number of studies have been undertaken (international and regional) ^1–5^ to provide context for this disconnect and to measure adult public opinions over time – with only marginal improvement in public opinion. This paper challenges the focus on adults and proffers an alternative focus on a significantly under-researched group: the early adolescent ^6^. With the abundance of data related to adult opinions about climate change, this paper examines the same (or similar) climate change opinions of 12-13-year-olds. We compare their opinions with their respective adult population, and also across two countries (Austria and Australia). This age group may provide a unique and unexplored avenue for climate science communication (Harker-Schuch, 2018); offering as-yet uncharted access to early worldview-construction development and, more critically, intellectual development pathways. In addition to improving understanding of climate science, such avenues may also improve support for climate-friendly policy and advocacy.

### The challenges of communicating climate change

Aligning public opinions with the scientific consensus on the influence of human-induced climate change is an ongoing challenge for both science communicators and those who recognise the essential role the general public play in mitigating and adapting to anthropocentric climate change ^8–10^. This challenge arises, in relation to science communication, on account of 1) individual socio-cultural/-political worldviews ^11^, 2) misinformation ^10,12,13^, 3) the cultivation of unwelcome emotions (anxiety, doubt, mistrust, confusion, overwhelming) ^14–16^ 4) the complexity and nature of the science ^17,18^, 5), a lack of common or shared knowledge, 6) the wicked nature of climate change and, lastly, 7) a lack of certainty ^19,20^ with regard to the scientific consensus or trust in the findings from the scientific community.

Navigating 1) worldview is, perhaps, the most pernicious challenge to overcome as it involves the individual’s idiosyncratic belief system and their attachment to social, political and cultural networks ^11,21^. Worldview is a factor known to not only prevent, but to inoculate an individual from re-assessing existing opinions or idiosyncratic belief systems ^22,23^. Attempts to promote re-assessment of beliefs can lead to an individual entrenching their opinion more firmly and undermining further communication efforts ^2^.

In addition to worldview, a great deal of 2) misinformation is generated both when lay-people attempt to rationalise and organise the information they are given or search for ^24^, as well as misinformation that is created by groups or industries that are threatened by a well-informed general public ^10,25–27^

With regard to 3) emotions, the issue of climate change has been found to engender fear ^28–30^, mistrust and doubt ^31^ (both with the scientific community and the broader body politic), as well as confusion about what to do and how to engage with climate change from a human behaviour perspective ^10,32^. Climate change is seen as an overwhelming problem and the mitigation/adaptation options available to individuals do not appear to offer adequate amelioration options – turning off the lights when you leave a room does not seem to be a meaningful response to the global threat of increasing atmospheric temperatures or rising sea levels ^33–35^. These emotional responses further paralyse effective engagement ^36^, polarise the issues associated with climate change and reinforce any existing worldview biases. The inability to effectively engage with climate change, coupled with unwelcome emotional responses, may also encourage individuals to ignore or down-play messages or information from science communicators as a means to limit further unwelcome emotions ^37^. As a species, we have never encountered a problem quite like the issue of climate change and we, therefore, lack the skills and perspectives required to analytically and methodically address it.

The 4) complexity of the science represents another barrier to science communication ^17,18^ as climate change involves many scientific disciplines (chemistry, physics, biology, geology) and is best understood with the expertise of highly specialised climate science fields within those scientific disciplines (e.g. chemistry: trace gases in Earth’s atmosphere, physics: astrophysics, biology: coral-mass accretion rates, geology: paleoclimatology and sediment accretion; to name but a very few) ^17,30,38^. Understanding the science is further hindered by the spatial and temporal scale of climate change which operates on spatial scales from the stellar to the molecular (which is unheard of in any other scientific discipline) as well as across time scales ranging from millennia to the momentary ^39,40^.

Because climate change wasn’t widely understood in the public arena until the mid-1980s ^41^, climate science was introduced to the curriculum quite recently and is, for most adults, a new science discipline. Very few concepts or ideas related to climate science were taught in school during their childhood and youth; creating a vacuum of 5) common, shared, or familiar knowledge. In addition, climate change is not taught well in most classrooms in the developed world ^12,42–44^ and/or is not necessarily offered as a part of the curriculum ^45^. To complicate matters further, climate change is considered a 6) ‘post-normal science’ or ‘wicked problem’ that is defined by criteria that make an effective solution (or compromise) difficult or impossible ^46^. Climate change is a wicked problem because it has no stopping rule, is both good and bad, is unique and unknown, could be seen as a symptom (as well as a cause), can have findings misinterpreted, has no – as such – definitive formulation, has innumerable solutions which cannot be immediately or ultimately tested, and failed solutions cannot be retracted and will not (and cannot) be tolerated ^47–49^. Lastly, although the ‘knowledge deficit’ model (being sufficiently informed on an issue i.e. climate science, in order to make an informed decision) has been largely dismissed ^50^ in view of worldview influences, the importance of climate literacy still forms a fundamental framework for opinion and engagement ^38,51,52^.

Finally, the lack of 7) confidence in the scientific community by members of the general public ^53^ (largely driven by targeted campaigns to promote mistrust or to misrepresent climate change as a ‘debate’) ^54^ has impeded climate science communication attempts – and severely retarded efforts to mitigate and adapt to climate change (particularly in the political arena). The single biggest influence on this impediment has been the notion of a ‘debate’ regarding climate science ^55,56^. Coupled with well-funded denialist campaigns, journalistic integrity to cover ‘both sides’ of an argument (or issue) have seriously interfered with establishing a well-informed public on the issue of climate change ^4,13^. Like the ‘flat-Earth’ concept, there is no viable counter- or alternative-argument to the theories that report the greenhouse effect phenomena – rendering the “false balance” reporting rationale of ‘both sides’ untenable ^57^.

The general scientific literacy of the public is variable, with even fundamental scientific principles poorly understood (e.g. the nature of the Earth’s orbit around the sun) ^58^, let alone the very specific scientific domains (as mentioned above: trace gas atmospheric chemistry - and so on) that form the scientific basis of climate change – leaving many lay-people unaware of the extraordinarily robust body of data (correlations, models and projections) and overwhelming evidence (proxy to empirical) of a human-induced climatic change. Nor is there a strong understanding in the public domain of the scientific consensus in relation to this issue– with 97% of climate scientists providing evidence of anthropogenic climate change ^59–61^. A recent study by the Pew Research Center in the USA (Politics of Climate Change, 2016), for example, indicated that although 50% of Liberal Democrats believed there was a scientific consensus on climate change, only 16% of conservative Republicans were of the same opinion – representing the strong gap between the public perceptions of, and the real, scientific consensus.

With these challenges in mind, and the continuing difficulties the general public have with regard to climate change policy and advocacy, we propose a new target for climate change interventions; one that may possess a stronger alignment with the scientific consensus and, as an extension of this alignment, be more receptive to interventions that promote climate-friendly behaviour and engagement – early adolescents.

### Why early adolescents matter

The early adolescent age group is important – they are the largest group of climate-vulnerable people on Earth and the group with the biggest portion of responsibility ^6,63^ – and they possess three vital characteristics that play a major role in an individual’s ability to comprehend the foundations of the climate change issue ^7,64–66^. The first is that their brains are undergoing a new intellectual development phase ^64,66^, the second is that their worldview has only just begun to form ^67,68^ and the third is a budding self-determination ^64–66,69^ that will, eventually, drive both their socio-political identity as well as help them secure social capital and community. These three characteristics, known as the 2^nd^ critical phase of development, arise as a result of physiological changes in the human brain that begin shortly before the age of 12 to ensure that healthy individuals will develop the skills they need to enter and manage adult life ^64^.

The intellectual development that takes place in this age group allows students to begin to process higher-order executive functions ^66^ and abstract reasoning processes. The mechanisms and processes that underlie climate change – particularly its ‘wickedness’ – require an individual to intellectually perceive the scale and connectedness of those processes and mechanisms. These perceptions are usually only possible once the brain begins this developmental phase ^64–66^.

As well as triggering executive function processing, the brain begins to form socio-political/-cultural worldviews ^67,68^. In conjunction with the abstract reasoning process, a proto-self-determination arises which is necessary for worldview development – making this age group an ideal ‘starting point’ for informed worldview development ^70,71^. The construction of these worldviews is strongly influenced by familiar others and by collective actions, attitudes or behaviours that are seen as desirable and help to secure an individual to their peers and respected others ^66^. There is a very short window of opportunity in this age group as worldview rapidly cements into attitudes of idiosyncratic belief systems that are, as has been previously explained, extremely difficult to alter ^38,46,68^. Recent research also indicates that embedding critical reasoning as an antidote to worldview-amendment resistance may offer an effective pathway for individuals to later re-assess their worldviews ^52^.

Lastly, as adolescents mature, they transfer their respect and social dependence from their parents and elders to their peers and familiar others ^65^. This transfer is a hallmark of adolescence and represents a significant social shift for science communicators both in terms of ‘potential for change’ that can be executed by that age group as well as a coalescence of new social networks (ibid). Young people tend to have high levels of social activism and this activism can lead to significant change throughout all levels of society ^72^. Aside from radical social adjustments, young people also implement gradual change as they secure relationships, find employment, enrol to vote and exercise their rights as adults through what they buy, who they associate with and how they manage their lives ^73,74^. Good governance depends, in many ways, on ensuring our youth are successfully transferred from childhood to adulthood and are able to find a place for themselves in society. Teaching them about climate change – both as a science and a wicked problem – will ensure they are prepared to engage with it successfully, and could also drive much-needed social coalescence on this issue ^56^.

This paper explores the suitability of this age group for science communication interventions toward improving their understanding and preparedness for the future. To this end, we attempt to determine the opinion signals of this age group in central urban centres both in comparison to their adult populations as well as to the scientific consensus. We also endeavour to determine how worried our early adolescents are with the issue of climate change as a measure of public concern. The tasks of adolescence are, of themselves, quite daunting and anxiety-ridden without the pressure and uncertainty of climate change ^65^. As adults, it is our duty to prepare young people for adulthood and to listen to their anxieties as a measure of their over-all emotional and mental well-being. Assessing 12 to 13-year-olds on their opinions related to climate change, therefore, becomes quite meaningful in broader social terms and may provide, for science communication professionals, a heretofore unexplored pathway for climate science communication.

### Objectives and Hypotheses

The overarching objective of this study is to determine the current opinion state of 12 to 13-year-olds with regard to the three arenas of worry (worry: is it something to worry about), the cause (human: is it anthropogenic), and its imminence (now: is it happening now) in relation to climate change and how these opinions influence one another (H1), and differ across country, school, gender, and preference of discipline (H2). Further, we review adult opinions in relation to climate change to investigate differences between the early adolescent and their respective adult population (H3). Lastly, we discuss the alignment of early adolescent opinions (worry, human and now) with the scientific consensus on climate change as a potential target for science communication interventions.

As such, this study explores the following hypotheses:

- H1: There is an influence of the different opinions (worry, human and now) on one another *(H0: There is no influence of the respective opinions on one another)*
- H2: The opinion of early adolescents on climate change differ based on demographic factors, such as country, gender, school, and disciplinary preference. *(H0: There is no difference in the opinion of early adolescents based on demographic factors.)*
- H3: The opinion of early adolescents on climate change is different from [or higher than] the opinion of adults in the same country. *(H0: There is no difference between the opinion of early adolescents and adults in the same country.)*

## Opinions – a synthesis of what adults currently believe

Before examining new data collected on the climate change opinions of early adolescents (12 to 13-year-olds), we first outline the current state and understanding of adult public opinion on climate change. While this paper is interested in, and reports on new data about, the opinions of the early adolescent, 12 to 13-year-old, age group, there is limited existing data on the opinions of this group (likely due to the impracticalities and challenges of research ethics of surveying children’s opinions) ^6,75,76^. As such, we collated the opinions of adults for the same themes (worry, cause, and imminence of climate change: ‘worry’, ‘human’ and ‘now’) from large opinion studies in order to compare adults to early adolescents as a measure of contrast; particularly in relation to worldview. Since adult opinions relating to climate change are well-studied and typically remain stable over time, they provide a useful group for comparison – not least if anticipated influences of adults over early adolescents (and their worldview) are observed to be less powerful than we may expect.

Opinions of adults on climate change vary from nation to nation ^77,78^ – and fluctuate within those nations over time, also ^20,79^. Determining the opinion of adults in relation to climate change allows science communicators to assess communication strategies and examine internal and external influences on opinions within communities (local to global) and monitor changing attitudes and perceptions. This process refines the science communication discipline and offers worthwhile insights into improving strategies for climate change as well as implementing strategies in other areas or in other science-related issues (e.g. GMOs, vaccinations). Although many surveys have been undertaken to monitor opinion in relation to climate change, there are differences in how they are constructed and how respondents are recruited. One difference is whether the response is yes/no or scaled on a preferential Likert scale (e.g. ranging between strongly agree to strongly disagree) (Leviston, Price, & Bishop, 2014). Such measurement differences make comparisons of results between surveys difficult. We can, however, see signals in those results that serve to inform science communicators and strengthen communication strategies within those respective countries. For some surveys, the data collection is across nations and national comparisons are made easier. Unfortunately, although opinion data has been collected over time from many nations, some countries lack sufficient data on the issue to gain more than cursory insights.

To establish a framework on which we can compare early adolescent opinions related to climate change, we looked at adult opinion data from the countries where we collected early adolescent data: Australia and Austria (as part of a larger research project investigating serious gaming interventions). Overall, Australians show that there is an influence of worldview with regard to climate change; political affiliation is a predictor of climate-related opinion ^55,81^ and there is a strong sceptic faction in Australia ^82^. While Australia has a plethora of literature on adult’s opinions with large-scale surveys going back to the early 1990s ^83,84^ literature on public opinions related to climate change in Austria, according to Rhomberg, ‘*is scarce and often focuses on Alpine regions*’ ^85^. Although this might be the case, Austria does show a very strong concern for issues related to climate change with 70% perceiving climate change as the world’s most significant problem (European Commission, 2014) and, following a recent EU-wide poll, 68% of Austrians consider climate change a ‘very serious problem’ ^5^. This concern reflects the amplified warming that Austria is currently experiencing in comparison to most other European countries ^87^ but it does not provide much insight into the nuances of public opinion, such as age, gender, education or relevant socio-cultural influences. Since the opinion signal from Austria is not as robust as from other EU nations, we have included opinions from France, Germany, Italy, the UK and Norway as a proxy measure in lieu of more specific opinion data from Austria and to obtain some understanding of European opinions as a whole (particularly as they are all long-term members of the European Union, are politically-significant EU members and/or are situated very close to Austria).

Differences among the countries of Europe are quite well-researched and -understood. France Norway, the UK and Germany have very strong support for opinion related to climate change ^1,88^ with Italy also demonstrating relatively high concern, albeit behind current economic concerns ^89,90^. In France, all major candidates in the last French presidential election supported efforts to mitigate and adapt to climate change. Norway, although a high emission-per-capita nation, lacks the polarized nature that is evident in Australia and the US – and strongly identifies as climate-friendly ^91^. The UK, although having a lower political impetus to address climate issues than other nations, has a strong environmental identity at the individual level. Germany, as an early-advocate for emission reduction ^41^, is a world leader in political efforts to develop the climate accord and, according to Schäfer, the ‘*dismissive segment*’*, which in the United States and Australia most strongly believes that climate change is not occurring or not caused by humans, is nonexistent in Germany%* ^88^. Opinion data from the United States has also been included for comparison of the EU data with a non-EU Western nation.

Opinion surveys have tended to measure the extent to which the public is worried or concerned, whether climate change is caused by human actions, and whether the climate is already changing. Data across these three dimensions are presented in the following sections.

### Are people worried about climate change?

#### Adult opinions – worry, now and human

For the opinion on whether climate change is something to worry about, the European countries all show stronger positive alignment with this opinion than Australia or the US (Chart 1a). For correlation to opinion topics (worry=worry, cause=human, imminence=now), please see Appendix 1. The opinion on the anthropogenic nature of climate change is quite different across the countries (Chart1b). Australia indicates a stronger opinion toward the influence of human-induced climate-change than the US but, again, the European countries all show much higher consensus with this opinion. For the opinion on whether climate change is happening now (Chart 1c), a greater proportion of Australians believe it is happening now than their US counterparts – although the proportion of people in Europe holding this opinion is higher still. Overall, we see that opinions in Europe are generally more aligned with evidence for climate change provided by the scientific consensus compared to Australia and the United States (which is generally lower again than Australia). For tables of adult data in relation to opinion topics (worry=worry, cause=human, imminence=now), please see Appendices 1 and 2.

**Figure 1:**
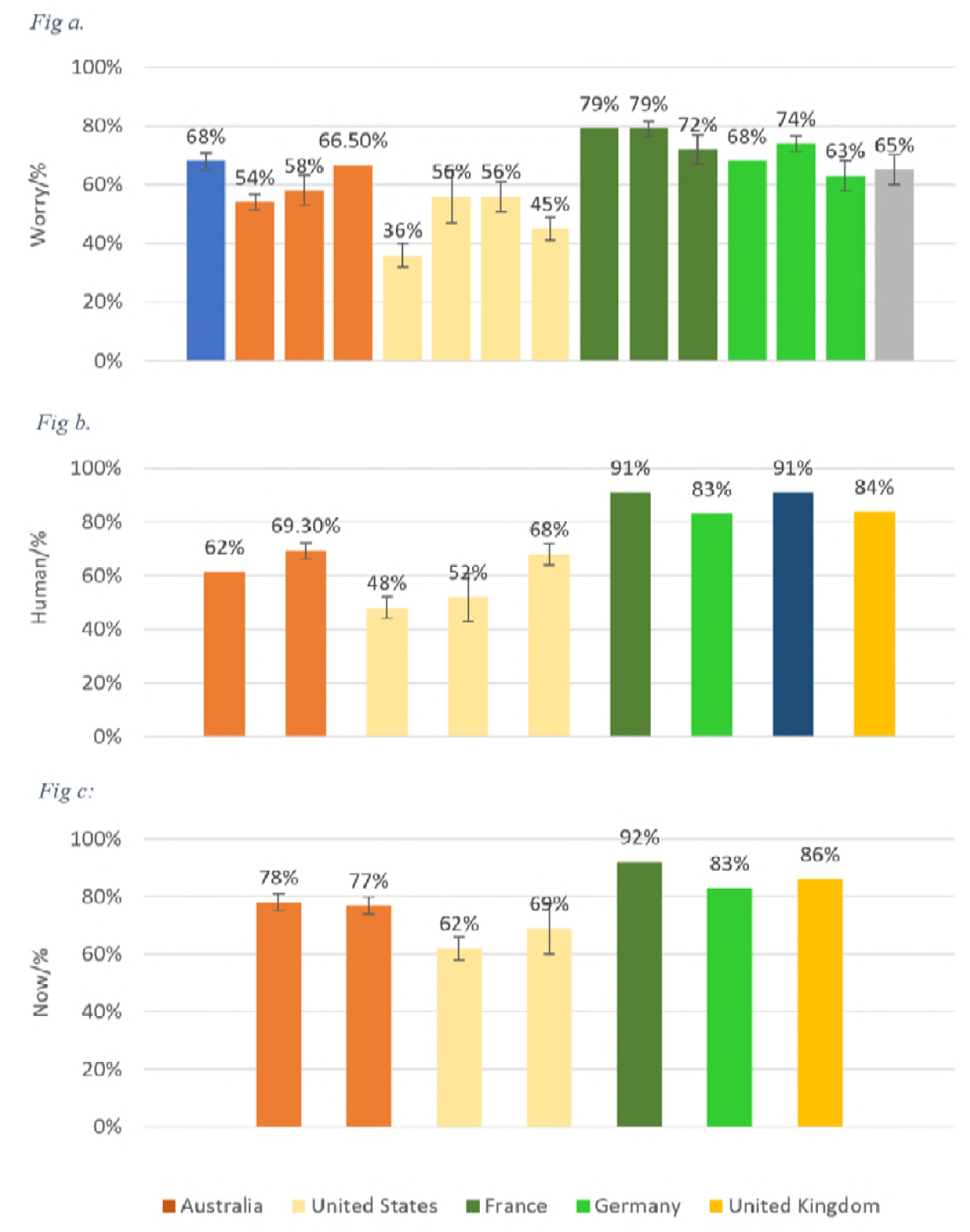
Collation of adult opinions in developed countries on whether (a) climate change is something to worry about (b) climate change is caused by human and (c) climate change is happening now. For data sources see Appendix 2. Error bars indicating uncertainty intervals as reported by the data sources have been included when available. The year in which each survey was conducted is shown below each survey result. For further details see Appendices 1 and 2.

## Materials and methods

To test the hypotheses, an opinion survey was created based on a previous survey by the primary researcher ^38^ and administered to first-year secondary students at six inner-urban high schools in Austria (in February-March, 2017) and Australia (June-August, 2017). This opinion survey, as previously mentioned, was part of a larger research project examining the role of serious-gaming interventions to improve climate literacy in the 12 to 13-year-age group (ethics protocol number: 2015/583). The survey was administered within the scheduled science class time (45-50 minutes) and was approved, as per the requirements for this research project, both by the relevant education departments and the ethics committee at the Australian National University. All protocols were followed in accordance with the requirements (ethical approval, anonymisation of the data, certifications for working with children/vulnerable people were met, and obtained permissions were stringently vetted: removing any participants where permission was not obtained).

### Schools and Students

The research catchment criteria for this study depended on schools being within a <10km driving proximity to the central business district and their status as secondary public school in an inner-urban setting. The selection of the school depended, as a requirement from the respective departments of education in Australia and Austria, on whether the school director and head of science would be willing to participate in this research. According to the requirements and procedures, 6 schools agreed to participate in this study (2 in Vienna, Austria - Coded as VHS1 and VHS2 - and 4 in Australia: 2 in Sydney - coded as SHS1 and SHS2 - and 2 in Canberra - coded as CHS1 and CHS2). All schools taught in the ‘mother-tongue’ of their respective nationalities (i.e. German in the Austrian schools and English in the Australian schools) and follow the state-regulated curriculum of their respective education departments.

The students were 12-13 years old and all first-year secondary students. A total of 834 students took part in the survey with a final 457 (207 (45.3%) females, 245 (53.6%) males, and 5 (1.1%) other) being eligible for final inclusion and analysis. Eligibility depended on approval from the department of Education, and the school, as well as parental and student approval, then participation in the study and valid responses to the survey.

Because the schools were ‘state suburb’ zoned (5 schools of 6) for their district or suburb (with one school allowing exceptional students to enrol) ^92^, we are not able to provide precise demographic information (census data) due to privacy laws as this is likely to make identification of the participating schools possible. It is, however, useful to provide some background information ^92,93^ and to note a few aspects of the demographics that may assist in interpreting the findings without compromising the privacy laws. To begin with, population density is higher in Vienna (176/ha) than in Sydney (27.6/ha) or Canberra (15.9/ha). CHS1 and SHS2 both had fewer citizens in the selected age group (12-13 years). Austrian citizens had a higher proportion of adults with only minimum-requirement education (age approx. 16). CHS1, VHS1 and VHS2 all had far higher non-mandatory secondary levels of attainment (approx. 18 years). CHS1, CHS2, SHS1 all had significantly higher tertiary levels of attainment (Bachelor and above). Canberra citizens have far higher ‘country of birth’ percentages than Sydney or Vienna and have significantly lower net immigration at present than the other schools. CHS1 had the lowest level of unemployment.

### Survey

In the first three items, the students were asked to put in an anonymous tracking code, their gender and preferred discipline. Following this, the next three items were Likert-style questions pertaining to their personal opinion with regard to their worry (is climate change something to worry about), the cause (is climate change caused by humans), and its imminence (is climate change happening now). The Likert scale ranged along a five-point scale:

No – Probably not – Maybe – Probably yes – Yes

For analyses, the Likert scale was converted to a numerical scale with No = 1, Probably not = 2, Maybe = 3, Probably yes = 4, and Yes = 5.

### Statistical Methods

Multiple regression models (IBM SPSS statistics 23.0) were used to analyse the respondents opinions about climate change. There were 3 variables, ‘Worry, ‘Human’, and ‘Now’ considered as the response variables (5-point Likert-scale as described above) as well the main effects of country, school, subject preference, and gender. Also, 2-way interactions for Country/School, Country/Subject Preference, Country/Gender were considered in the analyses. Coding for ‘Subject Preference’ explanatory variable (subjects from the respective schools: Austrian and Australia) were coded into their respective subject counter-part from the other school (i.e. ‘Science’ in Austria is separated into ‘Physics’, ‘Chemistry’, ‘Biology’ – all of these are distinctly different subjects in Austria but are taught under one subject ‘Science’ in Australia)) (Appendix 3).

The overarching analysis approach, therefore, consisted of the following stages:

1. Descriptive statistics on trends in overall opinion of early adolescents.
2. Multiple regression was conducted to determine the relationship between the response and the predictors

For the comparison on adult vs respondent opinions, and the between-country comparison, we calculated the comparisons of proportions using the ‘N-1’ Chi-squared test as recommended by Campbell (2007) and Richardson (2011). The confidence interval was calculated according to the recommended method given by Altman et al. (2000).

## Results

### Descriptive Data

#### Opinion data: climate change is something to worry about

In total, 302 students, corresponding to 66.1% of the students (*n* = 457), were firmly of the opinion that climate change is something to worry about (response = yes) and 102 students, corresponding to 22.3% of the students were of the opinion that climate change is something we should probably worry about (response=probably yes) (Table 5). The total positive response was 88.4%. The remaining responses (responses = ‘maybe’, ‘probably not’ and ‘no’) totalled 53 students and 11.7%, respectively.

**Table 1:** Overview of per school responses to the question: ‘In your opinion, is climate change something we should all worry about?’ Coding: VHS1 and VHS2: Vienna High School 1 and 2, respectively. SHS1 and SHS2: Sydney High School 1 and 2, respectively. CHS1 and CHS2: Canberra High School 1 and 2, respectively.

| Response | VHS1<br>n/ (%) | VHS2 n/<br>(%) | SHS1 n/<br>(%) | SHS2 n/<br>(%) | CHS1 n/<br>(%) | CHS2 n/<br>(%) |
| --- | --- | --- | --- | --- | --- | --- |
| Yes | 25<br>(58.1) | 22 (62.9) | 75 (60.0) | 52 (68.4) | 55 (67.9) | 73 (74.5) |
| Probably yes | 11<br>(25.6) | 8 (22.9) | 31 (24.8) | 16 (21.1) | 16 (19.8) | 20 (20.4) |
| Maybe | 6 (14.0) | 5 (14.3) | 13 (10.4) | 2 (2.6) | 7 (8.6) | 4 (4.1) |
| Probably not | 1 (2.3) | 0 (0.0) | 3 (2.4) | 4 (5.3) | 3 (3.7) | 0 (0.0) |
| No | 0 (0.0) | 0 (0.0) | 3 (2.4) | 1 (1.3) | 0 (0.0) | 1 (1.0) |
| Sum/country total:<br>'yes', 'probably yes' | 66 (85.3) |  | 338 (89.2) |  |  |  |

#### Opinion data: that climate change is caused by humans

In total, 253 students, corresponding to 55.4% of the students, were firmly of the opinion that climate change is anthropogenic in nature (response = yes) and 123 students, corresponding to 26.9% of the students were of the opinion that climate change is caused by humans (response=probably yes) (Table 6). The total positive response was 82.3%. The remaining responses (responses = ‘maybe’, ‘probably not’ and ‘no’) totalled 81 students and 17.7%, respectively.

**Table 2:** Overview of per school responses to the question: ‘In your opinion, do you think humans cause climate change‘? ‘ Coding: Same as for Table 1.

| Response | VHS1 n/<br>(%) | VHS2 n/<br>(%) | SHS1 n/<br>(%) | SHS2 n/<br>(%) | CHS1 n/<br>(%) | CHS2 n/<br>(%) |
| --- | --- | --- | --- | --- | --- | --- |
| Yes | 21 (48.8) | 16 (45.7) | 68 (54.4) | 42 (55.3) | 36 (44.4) | 70 (71.4) |
| Probably yes | 15 (34.9) | 5 (14.3) | 30 (24.0) | 21 (27.6) | 28 (34.6) | 24 (24.5) |
| Maybe | 2 (4.7) | 7 (20.0) | 18 (14.4) | 7 (9.2) | 11 (13.6) | 4 (4.1) |
| Probably not | 2 (4.7) | 2 (5.7) | 7 (5.6) | 1 (1.3) | 1 (1.2) | 0 (0.0) |
| No | 3 (7.0) | 5 (14.3) | 2 (1.6) | 4 (5.3) | 5 (6.2) | 0 (0.0) |
| Sum/country total:<br>'yes', 'probably yes' | 57 (71.9) |  | 319 (84.1) |  |  |  |

#### Opinion data: that climate change is happening now

In total, 268 students, corresponding to 58.6% of the students (*n* = 457), were firmly of the opinion that climate change is happening now (response = ‘yes’) and 124 students, corresponding to 27.1% of the students were of the opinion that climate change is probably happening now (response = ‘probably yes’) (Table 7). The total positive response was 85.7%. The remaining responses (responses = ‘maybe’, ‘probably not’ and ‘no’) totalled 65 students and 14.2%, respectively.

**Table 3:** Overview of per school responses to the question: ‘In your opinion, do you think climate change is happening now?’’ Coding: Same as for Table 1

| Response | VHS1 n/<br>(%) | VHS2 n/<br>(%) | SHS1 n/<br>(%) | SHS2 n/<br>(%) | CHS1 n/<br>(%) | CHS2 n/<br>(%) |
| --- | --- | --- | --- | --- | --- | --- |
| Yes | 17 (39.5) | 20 (57.1) | 77 (61.6) | 50 (65.8) | 43 (53.1) | 61 (62.2) |
| Probably yes | 15 (34.9) | 7 (20) | 36 (28.8) | 19 (25) | 22 (27.2) | 25 (25.5) |
| Maybe | 8 (18.6) | 7 (20) | 8 (6.4) | 6 (7.9) | 14 (17.3) | 8 (8.2) |
| Probably not | 0 (0.0) | 0 (0.0) | 2 (1.6) | 0 (0.0) | 0 (0.0) | 2 (2.0) |
| No | 3 (7.0) | 1 (3) | 2 (1.6) | 0 (0.0) | 2 (2.5) | 2 (2.0) |
| Sum/country total:<br>'yes', 'probably yes' | 59 (75.8) |  | 333 (87.3) |  |  |  |

### Statistical Analysis

#### Effect of opinions, country, school, gender and subject preference on opinions about climate change

The results of the multiple regression showed that all opinions were predictors of one another with statistically significant main effects of Human (χ^2^ = 47.98, df = 1, p-value = <.001) and Now ((χ^2^ = 14.12, df= 1, p-value = <.001). Both, Human (Model 2) (coef = 0.246, SE = 0.036) and Now (Model 3) (coef = 0.155, SE = 0.041) were positively related with the response Worry (Model 1). If a respondent had the opinion that climate change was something to worry about, this would likely predict that they also had the opinion that climate change was anthropogenic and was happening now (Table 8). Results for common explanatory variables are shown below for all response models.

**Table 4:** Response variable association with the statistical models

| Model | Response Variable | Explanatory variable |  |
| --- | --- | --- | --- |
|  |  | Opinion | Common |
| 1 | Worry | Now, Human | Country, School, Gender, Subject Preference |
| 2 | Human | Now, Worry |  |
| 3 | Now | Worry, Human |  |

**Table 5:** Comparison of 12-13 year old adolescents with respective adult population. @Data has been averaged from 2 or more surveys *P<0.0001 **P = P<0.01 ***no significant difference

|  | Austria |  |  | Australia |  |  | Between-country comparison |  |
| --- | --- | --- | --- | --- | --- | --- | --- | --- |
|  | Adolescents /% | Adults /% | Diff/ (adolescents – adults) /% | Adolescents /% | Adults /% | Diff/ (adolescents – adults) /% | Difference (Austrian – Australian) adolescents /% | Diff/ (Austrian – Australian) Adults /% |
| Worry | 86.1 | 71.32 | +13.98** | 89.1 | 63.28@ | +25.92* | -3.9*** | +8.04@* |
| Human | 72.9 | 87.24 | -11.44** | 82.1 | 63.69@ | +23.61* | -11.5** | +23.55@* |
| Now | 75.7 | 87.00@ | -15.34* | 87.6 | 77.72@ | +6.38** | -12.2** | +9.28@* |

### Model 1: Common effects on opinion that climate change is something to worry about

The results of the statistical analysis using multiple regression showed no significant main effect or 2-way interactions with any of the explanatory variables: country (χ^2^ = 0.186, df = 1, p-value = 0.667), school (χ^2^ = 10.248, df = 5, p-value = 0.69), gender (χ^2^ = 0.011, df = 1, p-value = 0.916), or subject preference (χ^2^ = 17.108, df = 10, p-value = 0.072).

### Model 2: Common effects on opinion that climate change is caused by humans

The results of the statistical analysis using multiple regression showed that although there is no difference in the main effect of opinion between the countries, per se, 2-way interaction between Country and Gender indicated that females in Austria (*n* = 33, mean = 4.33, SE = 0.162) are more likely than males in Austria (*n* = 48, mean = 3.93, SE = 0.140) to have the opinion that climate change is anthropogenic (χ^2^ = 9.530, df = 1, p-value = <.003). Additionally, males in Australia (*n* = 200, mean = 4.43, SE = 0.074) are more likely to have this opinion than females in Australia (*n* = 176, mean = 4.14, SE = 0.081; χ^2^ = 9.530, df = 1, p-value =<.003). Although the differences between means (i.e. 4.33 v. 3.93 and 4.43 v. 4.14,) in context of the Likert-scale, would be rounded to an integer of 4 (correlating to ‘probably yes’) and may seem marginal on the measurement scale, it is worthy to note that this prediction signal represents a difference between these groups of 8.0% (more female Austrians than male Austrians) and 5.8% (more male Australians than female Australians), respectively.

Subject Preference was also a predictor (χ^2^ = 18.475, df = 10, p-value = <.047), in that students who preferred certain subjects would more likely have the opinion that climate change was caused by humans with students who preferred Music (*n* = 9, mean = 4.65, SE = 0.288), Art (*n* = 47, mean = 4.50, SE = 0.143), Science (*n* = 59, mean = 4.35, SE = 0.124) or Sport (*n* = 119, mean = 4.19, SE = 0.090) more likely to have this opinion than those who preferred Technology (*n* = 19, mean = 3.93, SE = 0.214), History (*n* = 20, mean = 3.81, SE = 0.207). As with the above, although the actual difference between the means for each of these subject preferences does not appear profound when viewed in the context of the Likert-type scale (lowest subject rating 3.81 and highest subject rating 4.65) this difference, when applied to the sample population, represents a 16.8% difference (i.e. 16.8% more music, art, science and sport students think that climate change is anthropogenic than technology and history students).

### Model 3: Common effects on opinion that climate change is happening now

The results of the statistical analysis using multiple regression showed that 12 to 13-year-old Austrians (*n* = 83, χ^2^ = 4.375, df = 1, p-value = <.036; mean=4.06, SE = 0.150) were somewhat less likely to have the opinion that climate change is happening now than their 12 to 13-year-old Australian peers (*n* = 380, χ^2^ = 4.375, df = 1, p-value = <.036; mean=4.41, SE = 0.057). In addition, there was a 2-way interaction between Country and Subject Preference (χ^2^ = 18.514, df = 9, p-value = <.030), with students from Austria who preferred Mother Tongue (*n* = 1, mean = 2.09, SE = 0.812) and Music (*n* = 2, mean = 3.27, SE = 0.565) to be less likely to think that climate change was happening now (Chart 1) than their Australian peers who preferred Mother Tongue (*n* = 66, mean = 4.57, SE = 0.098) and Music (*n* = 8, mean = 4.47, SE = 0.283). In this case, when viewed in context with the Likert scale, the means range from 2.09 through to 4.47 and reflects a range difference of 47.6% (i.e. 47.6% more Austrian mother-tongue or music students were of the opinion that climate change is happening now than their Australian peers). This last interaction, Country with Subject Preference, can be explained by the low value for *n* in these categories.

**Figure 2:**
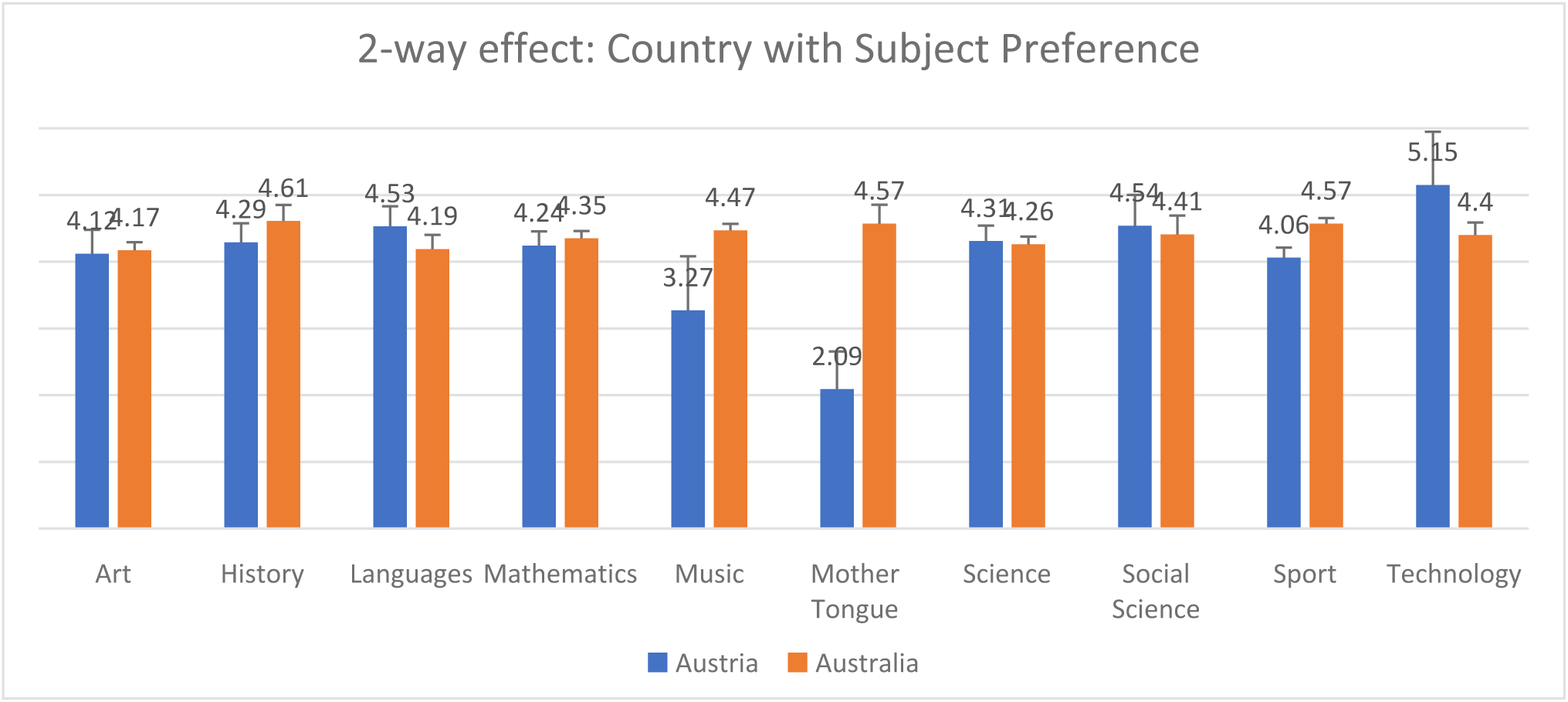
Subject preference effects on opinion that climate change is happening now. Covariates appearing in the model are fixed at the following values: Worry=4.51; Human=4.27. Please note that the estimated marginal means are based on the model, therefore, it could be possible that the mean value for country*Subject_preference is bigger than maximum value (e.g. Technology=5.164) i.e. Now = intercept + country + Subject_pref + country*Subject_pref + worry*B(worry) + Human*B(human) = 2.753 + 0.128(for country 1) + 0.6240(country(1)*Subject_pref(8)) + 0.194*(4.51 (fixed value)) + 0.184*(4.27(fixed value)) = 5.164

### Comparison of early adolescents with respective or proxy adult population

The following table provides an overview of early adolescent opinions in comparison to their respective (or proxy) adult population. Both student groups in Australia and Austria show a strong alignment with one another, a stronger positive worry level than Australian (63.28 %) and European (71.32 %;) adults which strongly supports the scientific consensus in the Worry opinions related to climate change. With regard to the Human and Now opinions, the Australian student group demonstrate a much higher level of concern than their Austrian peers and the European and Australian adults. In addition, more 12 to 13-year-olds in this study report the opinion that climate change is something to worry about (85.3 % Austrian respondents vs 71.32 % Austrian adults/89.2 % Australian respondents vs 63.29=8 % Australian adults) and this is further supported in the findings with students responding more definitely (yes= 63.83 %) to this opinion than to the Human (55.5 %) or the Now (56.33 %) opinion. Australian 12 to 13-year-olds in inner urban public-school environments do not reflect the same opinions for Worry and Human-causation as their respective adult population – our respondents were significantly more likely to think climate is something to worry about (89.2 % respondents vs 63.28 % adults) and is caused by humans (87.3 % respondents vs 63.69 % adults) with a lower difference for the opinion that climate change is happening now (84.1 % respondents vs 77.72% adults). In comparison, although Austrian 12 to 13-year-olds show a higher level of opinion for worry to their adult population (85.3% respondents vs 71.32% adults), they have significantly lower differences for the opinion Human (71.9% respondents vs 87.24% adults) and Now (75.8% vs 87.00%) as their respective European adult neighbours.

## Discussion

The study explored the opinion, and determinants, of 12 to 13-year-olds in relation to climate change, across the three arenas of worry, imminence, and human-causation. In light of the findings that each of the opinions (worry, human and now) predict one another, we reject the H1’s null hypothesis that there is no influence on the opinions for one another. The response for this age group in these areas indicates that the vast majority shares the concern that climate change is something to worry about, is caused by humans and is happening now –and these opinions relate predictably to one another.

With regard to the findings on the influence of demographic factors on opinion about climate change, we partially reject H2’s null hypothesis there is no difference in the opinion of early adolescents based on demographic factors, such as country, school, gender and subject preference. This is because some demographic factors correspond with significant differences in opinion on climate change, while others do not.

Lastly, as part of a broader research question, we determined differences between early adolescents and adults in the same (or proxy) country. We found that more adolescents than adults in Australia are concerned about climate change, whereas the comparison was more variable in Austria (and proxy countries).

Overall, we have found that opinions in the 12-13 year age group show strong pro-climate sensitivities – and the vast majority think climate change is something to worry about, is caused by humans and is happening now. The relation of the opinions to one another, though not surprising, is an important finding as it may allow us to extrapolate the same relationship in studies that have looked at only one aspect of these opinions. For science communicators, however, the demographic influences that affect an individual’s opinion are important and may offer insights into unexplored interventions and communication strategies.

## Demographic factors and opinion

### Climate change is something to worry about

In the statistical analysis, we see that the opinion for Now and Human strongly increases the opinion for Worry – meaning that those students who think that climate change is happening now and caused by humans are far more likely to be worried about climate change. These results are important as this suggests that worry regulation and emotional support would be worthy interventions in this age group – particularly those that foster hope and concern ^56,94^ as these are associated with stronger climate change beliefs, increased engagement and life satisfaction ^28^. This also suggests, due to the association of worry with climate change, that interventions that focus on causes (teaching the physical science basis: mechanisms, processes and basic climate science) instead of consequences (highlighting the impacts; sea-level rise, increased temperatures, extreme weather events) may diminish negative emotions associated with threats ^51^ and allow individuals to engage with climate change in a safer and more certain cognitive space of ‘normal’ science (as opposed to the ‘post-normal’ or wicked aspects of climate change). Establishing the ‘normal’ aspect of climate science may provide an intellectual scaffold to engage with that wickedness intellectually at a later age. Finally, these results also strongly reinforce previous research on emotional reasoning and associated changes in early adolescence which indicate that this age group are beginning to use ‘objective’, abstract-reasoning information to perceive threat ^71,95^.

### Climate change is caused by humans

Surprisingly, although there was no signal in the statistical analysis for country with regard to the opinion for Human (meaning that it didn’t matter which country the student came from with regard to the opinion that climate change is caused by humans) this influence became significant when gender was included in the analysis. Although research shows that late adolescent and adult females are more likely to be pro-environmental than males ^96,97^, our study suggests that this is not always the case, with Australian males reporting, more than their female peers, the opinion that climate change is caused by humans. Austria’s results supported the results of many previous studies with females showing a stronger opinion that climate change is anthropogenic than males. Of course, other factors may play a role and the lower number of Austrian respondents might suggest a stronger signal within a smaller demographic (community influences that we are not aware of, for example: immigration, zoning, lower school funding/resources). These findings suggest that research and tailored interventions aimed at targeting gender may be useful in promoting a better understanding of climate change. For example, serious gaming with a climate science topic may provide gender-specific game-play that responds to known gender differences – or, more usefully, are derived from game analytics that interact at the individual-student level to tailor learning to the learner needs.

Lastly, students who preferred Music, Art, Science, and Sport were more likely to support the opinion that humans are causing climate change than students who preferred Technology and History. The preference for science makes some sense, although previous studies in adults have argued that scientific literacy may decrease an individual’s support for this opinion, if they are ideologically predisposed to reject implications of climate change, e.g. economic or social changes ^58,68^. However, other studies rejected these findings as they found that the reported scientific literacy was not climate-science-specific or the literacy was self-assessed and this is problematic as people require an understanding of a scientific field in order to make an informed choice or, in the case of self-assessment, often claim more understanding than they really have ^51,68,98^. In addition, Shi et al. (2015) suggest that scientific literacy may not be an adequate predictor of climate literacy as expert knowledge is domain-specific and may not encompass other, unique areas of expertise. The preference for Music and Art may also align with communities where pro-environmental attitudes are common and strongly supported and engaged with. The preference for Sport is quite surprising and, aside from a (perhaps) more-frequent interaction with nature i.e. outdoor sports, weather-dependent play, this finding requires more investigation as it may indicate a nature-deficit influence ^99^. The lower opinions for Technology and History are, again, surprising. A rationale for this might be found in further research or, as mentioned above, a larger sample group across a broader swathe of the 12 to 13-year-age group. Importantly, though, while these differences were statistically significant, all mean ratings from each subject area were situated between ‘maybe’ and ‘definitely yes’, so the effective difference in opinion, while noteworthy, does not indicate a profound difference in opinions based on subject preference.

### Climate change is happening now

The most surprising finding of this study is the stronger opinion amongst 12 to 13-year-old Australian public-school students living in central urban districts that climate change is happening now than is shared by their Austrian peers (84.1% Australian respondents vs 71.9% Austrian respondents). It is especially remarkable that, in light of the amplified warming that is taking place in Austria ^85,100^, that Austrian students are less likely to have the opinion that it is happening now. In addition, the 2-way interaction between Country and Subject Preference, Australian students who prefer Music and Mother Tongue, were significantly more likely to think that climate change is happening now than their Austrian peers who prefer these subjects, with the means for each subject area representing a spread from ‘probably not’ (Austria) to ‘probably yes’ (Australia).

### Comparison of adolescents with adults

Although we might anticipate a strong alignment with the respective political position on climate change in each country (i.e. strong positive adolescent and adult opinions in Austria in line with EU climate policy and weaker positive adolescent and adult opinions in Australia in line with weaker Australian climate policy), we found that Austrian student were less likely to have the opinion that climate change is happening now and is caused by humans – both in comparison to their proxy adult population (Human: 75.8% Austrian respondents vs 87.24% Austrian^1^ adults; Now: 71.9% Austrian respondents vs 87.0% Austrian^1^ adults) and to their Australian peers (Human: 75.8% Austrian respondents vs 87.3% Australian respondents; Now: 71.9% Austrian respondents vs 84.1%; Australian respondents). This finding challenges the anticipated influence of their adult populations – especially as the comparison shows Australian 12 to 13-year-olds think climate change is something to worry about, is caused by humans and is happening now, more than their adult cohort do (Worry: 89.2% respondents vs 63.28% adults; Human: 87.3% respondents vs 63.69% adults; Now: 84.1% respondents vs 77.72% adults) – although Austria 12-13 year olds are more worried than their respective adult population (Worry: 85.3% Austrian respondents vs 71.32% Austrian^1^ adults), they show lower opinion levels for Human (75.8% respondents vs 87.24% adults^1^) and Now (71.9% Austrian respondents vs 87.00% Austrian^1^ adults) than their proxy adult population.

There may be differences in culture or lifestyle between adolescents in Austria and Australia, such as differences in population density or interactions with nature ^101^ that lead to the observed differences in opinion. However, it would be likely to see any such effect reflected similarly in the adult populations if it is simply an effect of place. Instead, if there is no methodological or measurement error responsible for the difference, then these results indicate there is an interaction between the adolescent experience and place which shape the attitudes. For example, curriculum content or norms around adolescents’ awareness of climate change or other key policy issues. Curiously, both Australian and Austrian 12-13 year olds show higher rates of reporting the worry opinion when compared to their respective adult populations – and with a stronger positive response than for the other opinions (now and human). This worry signal is an important one as it suggests that, although Austrians in this age group are attuned to the emotional aspect of climate change as a threat, they do not possess the fundamental understanding of climate change processes to recognise the major dimensions of climate change which make it worthy of worry; these are both the imminence of the threat (now), and the fact that the observed warming and climatic changes are resulting from human interference in the climate system (human).

### Limitations to this study

It is necessary to note that certain biases may have influenced the data and affected the findings. The first is that the selected students were from a total of six schools, and as a result cannot be considered a geographically or demographically representative sample of either country. Despite this, the results are useful, especially as data on the 12 to 13-year-old age group is scarce in the literature. It would be beneficial for future studies focused on early adolescents to adopt compatible methods to allow for aggregation of data, developing a more robust data set. One of the barriers to more geographically and demographically representative data from 12 to 13-year-olds is the (necessary) challenge posed by research ethics of working with young and vulnerable people. All participants required approval from the school, their teachers, the parents, and the students themselves. Those who maintain climate-friendly attitudes are, therefore, more likely to participate in this research than those who do not. The level of teacher engagement was, perhaps, the most influential of all the potential biases for the teachers were the essential driver behind participation numbers in each class. The author observed that the teachers who were not enthusiastic had a far lower number of participants in their class than those who were favourable towards the research. This observation was apparent in anecdotal negative criticism of the project by those teachers who returned fewer participation notes from their students and, in some cases, suggesting to the researcher that climate science was not a ‘settled’ science. In addition, one of the schools in Vienna (VHS2) had parents that were very sceptical about their child’s involvement in a research project with 2 out of the 4 classes returning notes that denied permission. Many of these parents were new residents in Vienna (very recent arrivals), so it was difficult to discern whether declining to participate was on account of their vulnerability as new residents or due to negative attitudes toward climate change. If the latter, then these important perspectives were not able to be captured in the study. Curiously, nearly all permission notes were returned by the parents in the Austrian schools (even those stating that their child could not participate) whereas just over half were returned from Australian schools (with nearly all saying their child could participate) even though the recruitment process had been the same. The researcher speculates whether the unreturned notes in Australia are in lieu of a returned note that does not allow their child to participate or a lack of procedure between the school and home that results in lost or misplaced permission notes – or a mix of both. These unavoidable challenges of working with schools and their adolescent students are useful for other researchers to note when engaging with similar samples for future research.

### Implications of this study

As worldview plays such a significant role in the opinions and behaviour of adults, the 12-13-year age-group presents an opportunity for science communication intervention. While socio-cultural worldviews are still nascent, and not strongly influencing opinion development, the intellectual scaffolding for understanding climate science is far-more strongly developed at this age. The potential to improve an individual’s understanding of climate science as a basis for engaging with climate-friendly policy and advocacy, without the bias of worldview, makes this age group an ideal target. In the 12-13-year age group, addressing the ‘knowledge deficit’ (being sufficiently informed on an issue i.e. climate science, in order to make an informed decision) ^38,51,68^ is a valid pathway for interventions as these individuals are all, without exception, enrolled in school in order to be given the information, literacy, skills and intellectual tools they need to enter society.

Although climate science is complex, this age group has already begun to cultivate the cognitive framework that can allow them to intellectually process the different aspects of climate science – including feedbacks, interactions, and scales. This is especially evident in the increased sense of worry they show if they do think that climate change is happening and is caused by humans.

## Conclusion

The potential for the 12 to 13-year-old age group as important targets for science-based climate change education is clear. Not only do we have an age group whose opinions already align well with the scientific consensus, we also have a group that could greatly benefit from well-designed science communication interventions due to the stage of their intellectual development. Additionally, early adolescents are easy to reach as they are all in school, and they are at the nascent stage of worldview construction. Improving scientific literacy in relation to climate change could have immense social and political implications, such as providing all young people with a fundamental understanding of the science of climate change, regardless of the political ideology or social identity they will develop in the years ahead. Perhaps, if such a literacy programme was properly implemented, we would have a general public that, regardless of worldviews and belief systems, would share a good understanding of the science of climate change as the basis for public and policy deliberations on relevant courses of action. Climate-science education of early adolescents offers alternative intervention routes that avoid the worldview-based polarisation on the reality of climate change which we have experienced in recent decades. Future climate science-educated adults could no more deny the phenomena of climate change than they could deny the existence of their large intestines: both are physical phenomena manifest invisibly in our everyday lives.

## Acknowledgements

Dr Hwan-Jin Yoon, Statistical Consulting Unit, The Australian National University

Ms Michel Watson, CPAS, The Australian National University

This research is supported by an Australian Government Research Training Program (RTP) Scholarship.

## Footnotes

1 Where opinion data is not available for Austria proxy data has been used (e.g. neighbouring EU countries)

